# Estimating protein-ligand interactions with geometric deep learning and mixture density models

**DOI:** 10.1101/2023.10.03.560738

**Authors:** Yogesh Kalakoti, Swaraj Gawande, Durai Sundar

## Abstract

Understanding the interactions between a ligand and its molecular target is crucial in guiding the optimization of molecules for any *in-silico* drug-design workflow. Multiple experimental and computational methods have been developed to better understand these intermolecular interactions. With the availability of a large number of structural datasets, there is a need for developing statistical frameworks that improve upon existing physics-based solutions. Here, we report a method based on geometric deep learning that is capable of predicting the binding conformations of ligands to protein targets. A technique to generate graphical representations of protein was developed to exploit the topological and electrostatic properties of the binding region. The developed framework, based on graph neural networks, learns a statistical potential based on the distance likelihood, which is tailor-made for each ligand–target pair. This potential can be coupled with global optimization algorithms such as differential evolution to reproduce the experimental binding conformations of ligands. We show that the potential based on distance likelihood, described here, performs similarly or better than well-established scoring functions for docking and screening tasks. Overall, this method represents an example of how artificial intelligence can be used to improve structure-based drug design.

**Significance statement:** Current machine learning-based solutions to model protein-ligand interactions lack the level of interpretability that physics-based methods usually provide. Here, a workflow to embed protein binding surfaces as graphs was developed to serve as a viable data structure to be processed by geometric deep learning. The developed architecture based on mixture density models was employed to accurately estimate the position and conformation of the small molecule within the binding region. The likelihood-based scoring function was compared against existing physics-based alternatives, and significant performance improvements in terms of docking power, screening power and reverse screening power were observed. Taken together, the developed framework provides a platform for utilising geometric deep-learning models for interpretable prediction of protein-ligand interactions at a residue level.

## 1 Background

Identifying potent drug candidates from the vast chemical space (10^60^ drug-like molecules) is non-trivial, and their accurate identification is fundamental to the discovery, repurposing and development of lead molecules [1]. While biochemical experimentation is limited by the aspects of time and scale, *in-silico* methods have been reliant on physical representations of proteins and small molecules to generate mathematical models that mimic the behaviour of these biological entities. Methods such as X-ray diffraction, NMR crystallography and others have been providing an estimation of the 3-dimensional state of a given biological molecule with a reasonable degree of fidelity [2–4]. This information, in the form of machine-interpretable formats like (pdb, sdf) are utilized to model the behaviour of proteins in the biological system and also their interaction with other entities like small molecules. It is well established that a few key interactions between the ligand and its molecular target determine the efficacy, potency and selectivity of the given drug. Methods like molecular docking/dynamics allow us to understand these interactions and explore the chemical space for favourable interactions [5,6]. In recent years, with the advancements in experimental structural biology, there has been an influx of a tremendous amount of data. Although traditional physics-based methods were considered capable of processing large numbers of structures with virtual/reverse-virtual screening techniques, they are apparently being limited by the sheer amount of data being presented.

AI-assisted computational structural biology is quickly progressing in the direction of being competitive with the traditional *in-silico* methods that employ physics-based workflows to model the interaction between biological molecules [7,8]. While there are numerous attempts to develop statistics-based methods for scoring ligand-target combinations, there is minimal focus on getting optimum conformations of the ligand-target system in a machine-learning platform. Further, the lack of residue-level interpretability is one of the biggest factors that favor the usage of physics-based methods despite their limitations in terms of speed and scale. Generating efficient representation of data is an extremely vital aspect of building an efficient ML agent [2]. More so in the case of biological molecules, where the distribution of data is highly multivariate, and there has been a trend of using sequences and chemical descriptors for data representation [9,10]. In contrast to fields such as machine vision and voice recognition, generating reliable representations of molecular data is challenging. Conventionally, there has been a reliance on vector encodings and fingerprints to embed the molecular information into a matrix of values that can be further used to train ML models [11,12]. However, molecular data is fundamentally different from image or voice, and has a spatial aspect to them in addition to their physical and chemical characteristics. Geometric representation of molecules presents a platform to incorporate the topological arrangement of atoms in 3-dimensional space as a trainable data structure [13]. These graph-based representations, in tandem with pre-existing chemical and sequence embeddings (from pertained language models), can improve the overall reliability of the predictors. Graph neural networks (GNNs) have emerged as a powerful tool for representing and processing complex relational data [14]. GNNs offer a promising alternative by providing a way to encode the structural and relational information of molecules in a graph-based representation that can be easily processed by a neural network. They have been applied to tasks such as predicting the efficacy of potential drug molecules and identifying potential drug-drug interactions [9,15,16].

Here, we present *Alphadock*, a geometric deep learning-based framework for identifying interactions between a given drug-target pair at a residue level. *Alphadock* incorporates the topological and electrostatic characteristics of protein surfaces into the machine-learning framework as graphs. In addition to that, pre-trained protein language models were employed to extract sequence embeddings as a vital source of trainable information. Coupled with elements of geometric deep learning, the proposed framework was able to estimate ligand position and conformation at the protein binding site with a high degree of fidelity. Moreover, the developed distance-likelihood-based scoring function performed at par with existing physics-based scoring functions in multiple metrics of evaluation.

## 2 Results

Existing machine learning models to estimate drug target interactions generally model the likelihood of inter-nodal associations within protein and ligand based on the ground truth and feature sets [17]. While accurate ground truths in the form of interatomic distances between protein and ligands are available in curated datasets such as PDB, there is still a bottleneck in terms of extracting efficient representations from protein structures. Most of the studies have employed amino acid sequences as the primary source of information to train the machine learning model, and hence, leaving a lot on the table in terms of structural features. Here, structural features, in the form of graphs, along with sequence embeddings were utilized for training a geometric-deep learning model to estimate distance likelihood for a given ligand target pair with a high level of accuracy. Additionally, the proposed workflow could also determine the reliable conformation state of the ligand in the binding pocket with the use of an optimization algorithm.

### 2.1 Mixture density networks as an alternative to regression-based approaches for quantifying protein-ligand affinity

The core principle of MDNs lies in modeling the conditional distribution of the target variable given the input variables using a mixture model [18]. This involves representing the target distribution as a weighted sum of several component distributions, each governed by its own set of parameters. These parameters, such as means, variances, and mixing coefficients, are learned by the MDN through the training process. MDNs leverage the expressive power of neural networks to estimate these mixture model parameters. The input variables are passed through a neural network, which outputs the parameters of each component distribution.

The network is trained by optimizing a suitable loss function, typically based on maximum likelihood estimation, to maximize the likelihood of generating the observed target values given the input variables. The developed MDN network comprises a fully connected deep neural network that was trained to estimate the parameters of a gaussian mixture model such as means, standard deviations and mixing coefficients. On the contrary, a conventional approach employing regression would minimize a metric (mean absolute error or root mean squared error) based on the aggregate distances between the ligand and the target [19]. This would be inadequate to model the scope of interactions and also the interpretability of the trained models. **Table 1** compiles the performance of multiple variants of the proposed workflow in terms of metrics such as root mean squared deviation (RMSD), Pearson’s correlation, spearman correlation and Kendall rank. It can be observed that the RMSD of the estimated ligand conformation in the binding pocket is within the accepted limits of around 2 Å for most of the variants. Similarly, the workflow shows a decent statistical correlation between the scores of the predicted conformations and the native binding pose.

**Table 1:**
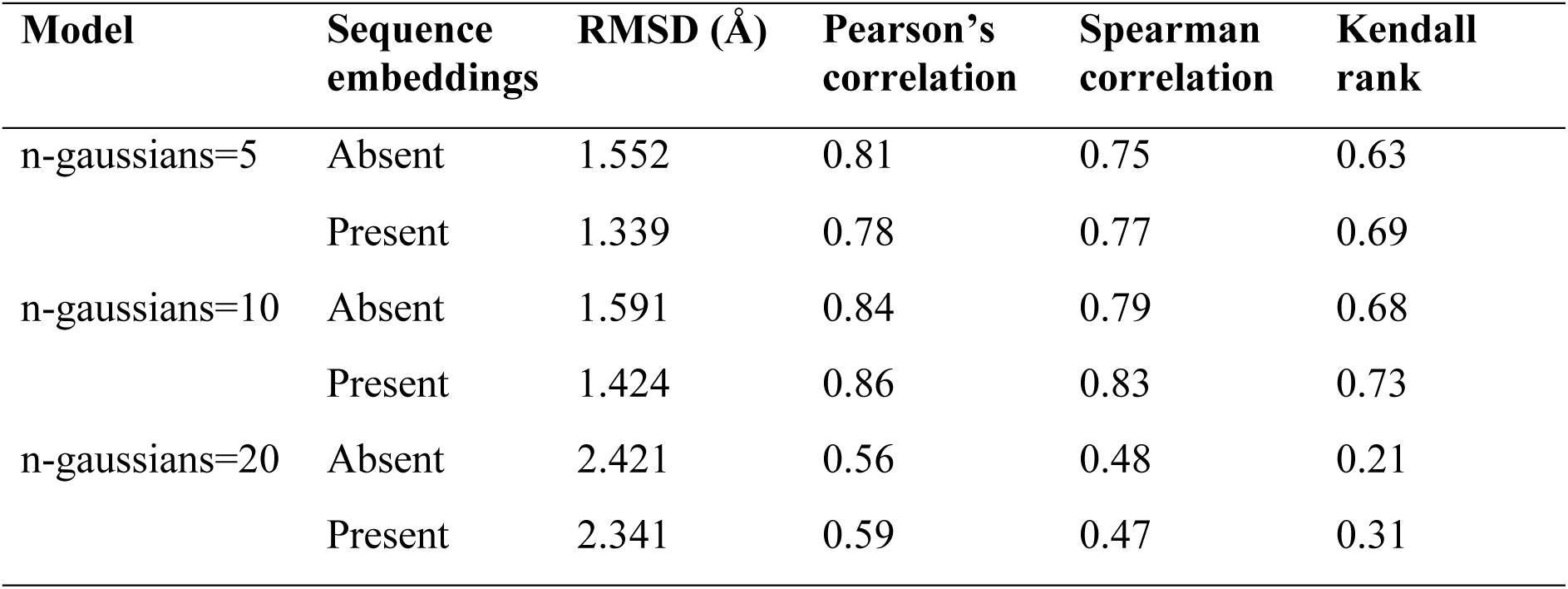
The effect of number of gaussians in the multi density network analysis for estimating the binding conformation.

| Model | Sequence embeddings | RMSD (Å) | Pearson’s correlation | Spearman correlation | Kendall rank |
| --- | --- | --- | --- | --- | --- |
| n-gaussians=5 | Absent | 1.552 | 0.81 | 0.75 | 0.63 |
|  | Present | 1.339 | 0.78 | 0.77 | 0.69 |
| n-gaussians=10 | Absent | 1.591 | 0.84 | 0.79 | 0.68 |
|  | Present | 1.424 | 0.86 | 0.83 | 0.73 |
| n-gaussians=20 | Absent | 2.421 | 0.56 | 0.48 | 0.21 |
|  | Present | 2.341 | 0.59 | 0.47 | 0.31 |

### 2.2 Including sequence embeddings along with structural features improves performance

Protein sequence embeddings are representations of protein sequences in a lower-dimensional space, capturing the underlying features and relationships between amino acids. By combining sequence embeddings with structural features, such as hand-engineered features or features extracted from domain-specific knowledge, a model can learn from a more comprehensive and diverse set of information [20]. By combining sequence embeddings and structural features, improved performance on the target task was expected, as the model would be able to make use of both the inherent patterns in the data and the domain-specific knowledge encoded in the structural features. A variant of the ESM model was employed to encode the protein sequence and fed it to the model along with the graph-structured data [21]. As expected, a marginal improvement of 8.43% (averaged over all variants) in terms of RMSD was observed on the validation set, as summarized in **Table 1**. This further validates the hypothesis that protein sequence embeddings from pre-trained language models contain vital information that can be employed as accessory features for improving performance.

### 2.3 The proposed statistics-based scoring function is comparable with existing physics-based scoring functions

Scoring functions are designed based on physical and empirical principles that govern molecular interactions. They attempt to quantify the intermolecular forces and energy terms involved in ligand-receptor binding, such as van der waals interactions, electrostatic interactions, hydrogen bonding, and solvation effects. The goal is to accurately rank and prioritize ligands based on their binding affinities, ensuring that ligands with higher predicted scores are more likely to bind strongly to the target protein. The effectiveness of a scoring function can be evaluated with estimates such as docking power, screening power, and scoring power. CASF-2016 benchmark datasets contain hundreds of decoy conformations for all the 285 protein-ligand complexes [22]. The standard deviation (RMSD) for these decoys ranged from 0-10 Å. An effective scoring function is expected to rank these decoys so that the conformations that are closest to the correct binding pose (< 2Å) are ranked at the top. The proposed method and its scoring function were compared with 35 commonly used scoring functions and **Figure 2a-b** summarizes the results for docking power, forward screening power and reverse screening power for CASF16 benchmark sets.

**Figure 1:**
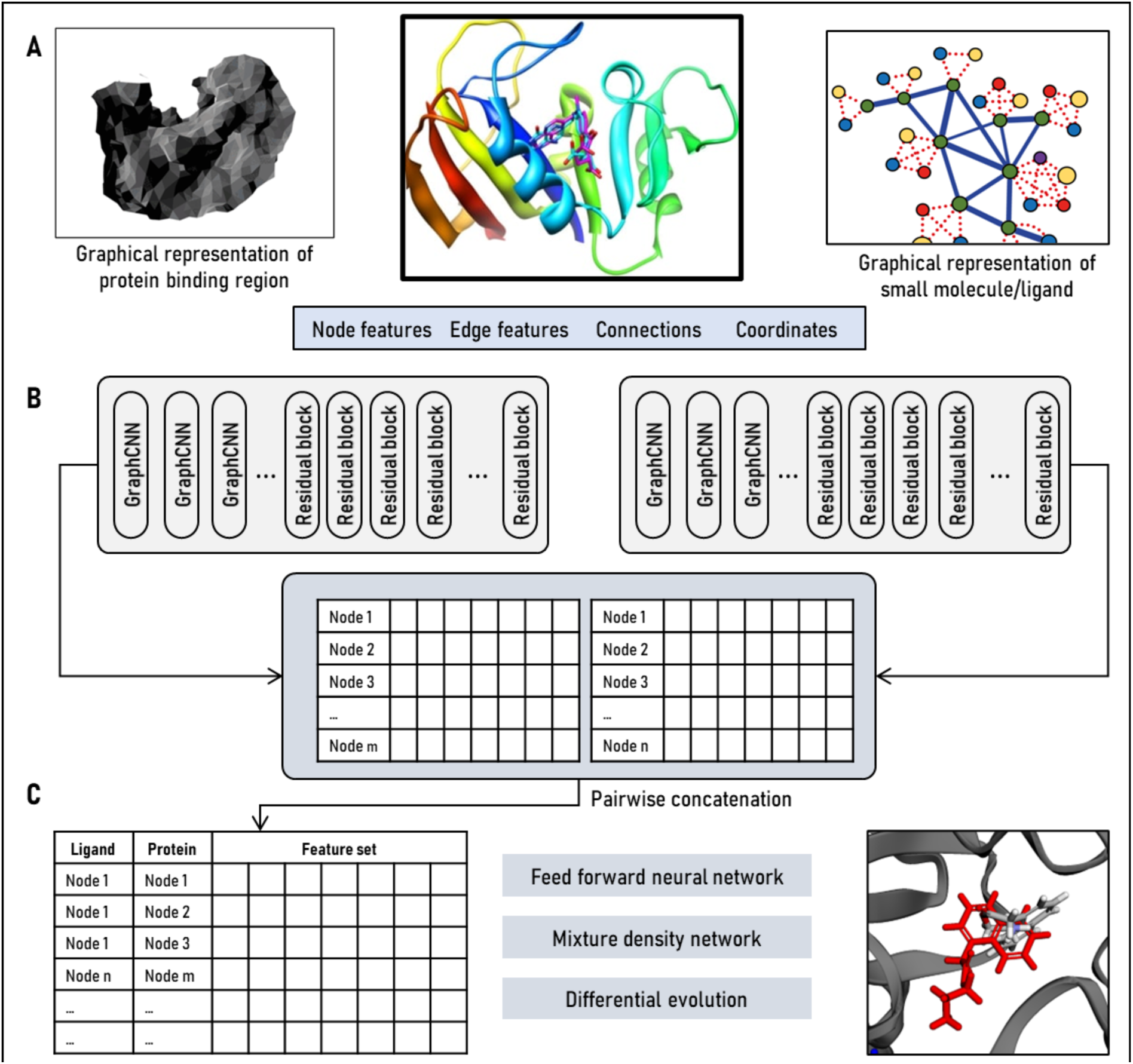
Overall structure of the developed *Alphadock* framework. (A) Protein and ligand are represented as their respective graphical forms. (B) gCNN and residual blocks to extract features from protein and ligand graphs followed by pairwise concatenation. (C) Processing of concatenated feature vectors through feed forward neural network, mixture density models and optimization algorithm to generate accurate ligand conformation.

**Figure 2:**
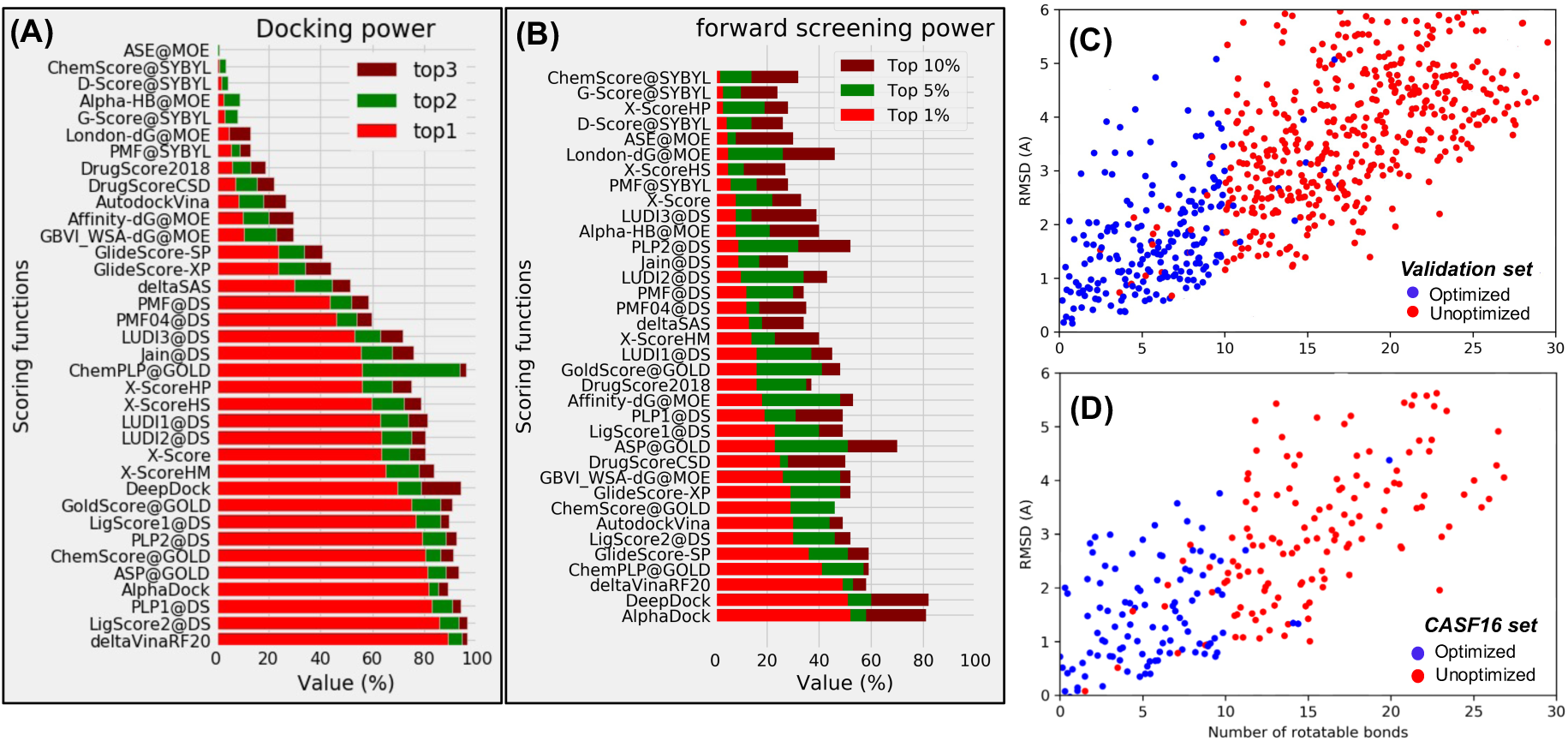
Comparative analysis of the *Alphadock* scoring function with already existing physics-based functions on the CASF 2016 benchmark dataset. The figure compiles results of comparative analysis for (A) Docking power and (B) Forward screening power. Dependence of number of rotatable bonds on RMSD of predicted conformation for (C) internal and (D) external validation (CASF16) set.

Docking power is the measure that defines the ability of the scoring function to correctly identify molecules that are closest to the natively binding ligand among synthesized decoys with similar conformation [23]. Similarly, screening power refers to the ability of the scoring function to effectively prioritize and identify potential binding interactions between a small molecule (ligand) and a target protein or receptor. A scoring function with high screening power will be able to successfully identify ligands that have strong binding affinity and are likely to exhibit desired biological activity. This involves accurately predicting the binding mode, energy, and affinity of the ligand-receptor complex. It was observed that the top-ranked decoy from our method was within the < 2Å RMSD threshold for 91% of the 285 protein-ligand complexes under evaluation.

Furthermore, it was observed that the number of rotatable bonds is a very important factor while running the molecular optimization protocol. As evident from **Figure 2c-d**, the optimizer is not able to process molecules with a higher number of rotatable bonds in internal as well as external validation sets, which is quantified by the RMSD scores.

### 2.4 Employing distance likelihoods in the scoring function aids in estimating molecular conformations with a high degree of confidence

The trained model was clubbed with an optimization algorithm to generate the ligand conformation with the highest likelihood of binding with the given protein. Distance likelihoods that were provided by the trained MDN model presented a statistical measure of the likelihood or probability of observing certain interatomic distances in a given molecular conformation. Out of the many optimization algorithms, such as differential evolution, simulated annealing, and genetic algorithms, differential evolution was chosen for its ease of implementation [24,25]. For the current use case, differential evolution worked by maintaining a population of candidate solutions, called “individuals,” and iteratively applying molecular operations (rotations and translations) to generate new conformations that are hopefully better than the previous ones [26]. The algorithm selected the best individuals from the population and used them to generate the next generation. This process was repeated until the desired optimization level or a predetermined number of iterations was reached. The methodology proved to be effective in generating accurate conformations of molecules in the validation set (**Figure 2**).

## 3 Discussion

*Alphadock* was developed to learn the potential and optimal binding conformation that is specific to each ligand-protein complex. It employed experimental 3D complexes of bound proteins and ligands retrieved from PDBbind database for training. The entire workflow of *Alphadock* training involved two steps, namely, (i) feature extraction and processing to encode the local environment of the ligand and protein, and (ii) making use of the encodings to estimate the relative location and interactions within protein and ligand accurately.

Ligand molecules were directly represented as undirected graphs with nodes and edges representing atoms and bond-types respectively. On the other hand, building graphs from protein binding surfaces was a bit complex. The protein binding surface was converted into a mesh-like structure (consisting of nodes, edges and faces) taking inspiration from pre-existing surface fingerprinting methods like Masif. The protein ‘mesh’ was encoded with electrochemical properties, connectivity information and 3D coordinates of individual nodes. In the first step of active feature processing, stacked residual GNNs were used to extract the local environment of protein and ligand from their graph representations. Employing GNNs to process features allowed the model to not only hold information about the individual nodes, but also have a sense of their surroundings. Although some pre-existing methods have tried to process graph representations of molecules as previously described, most of them follow the workflow with a feed-forward neural network that is trained by minimizing the error (mean squared, average distance). However, the relative distances between individual nodes in the protein and ligand graphs have a multimodal distribution owing to the differences in the degree of likelihood with which the individual protein-ligand nodes are separated from each other. Feed-forward neural networks trained on average statistics might not be very effective in this case. Therefore, multi-density networks (MDNs) were employed as a downstream architecture to account for the multi-modal nature of data. MDNs consist of a neural network with one or more output layers that are used to model the parameters of a mixture model. A mixture model is a probabilistic model that represents a distribution as a mixture of several component distributions. The component distributions in a mixture model can be any type of distribution (gaussian in this case), and the mixture model is defined by a set of weights that indicate the relative contribution of each component to the overall distribution. The results clearly outlined the effectiveness of employing MDNs for modelling distance-based likelihood estimates. Interestingly, as observed in **Table 1**, a model that was trained to estimate the parameters of a mixture model with ten gaussians performed better than the mixture models with five or twenty gaussians. Therefore, it can be inferred that merely increasing the number of gaussians for modelling distance likelihoods does not guarantee an increase in model performance. However, the effect of including sequence embeddings in the training protocol was clearly visible in the same results.

Also, while optimizing the given ligand molecule for best 3D conformation, it was interesting to note that the number of rotatable bonds was a primary factor (**Table 2**). This was quantified by the lower RMSD value for the molecules with lesser rotatable bonds. A variation in model performance was observed for different enzyme types in the validation set, as depicted in **Figure 3**. It can be explained on the basis of variation in the general structure and function of these enzymes.

**Figure 3:**
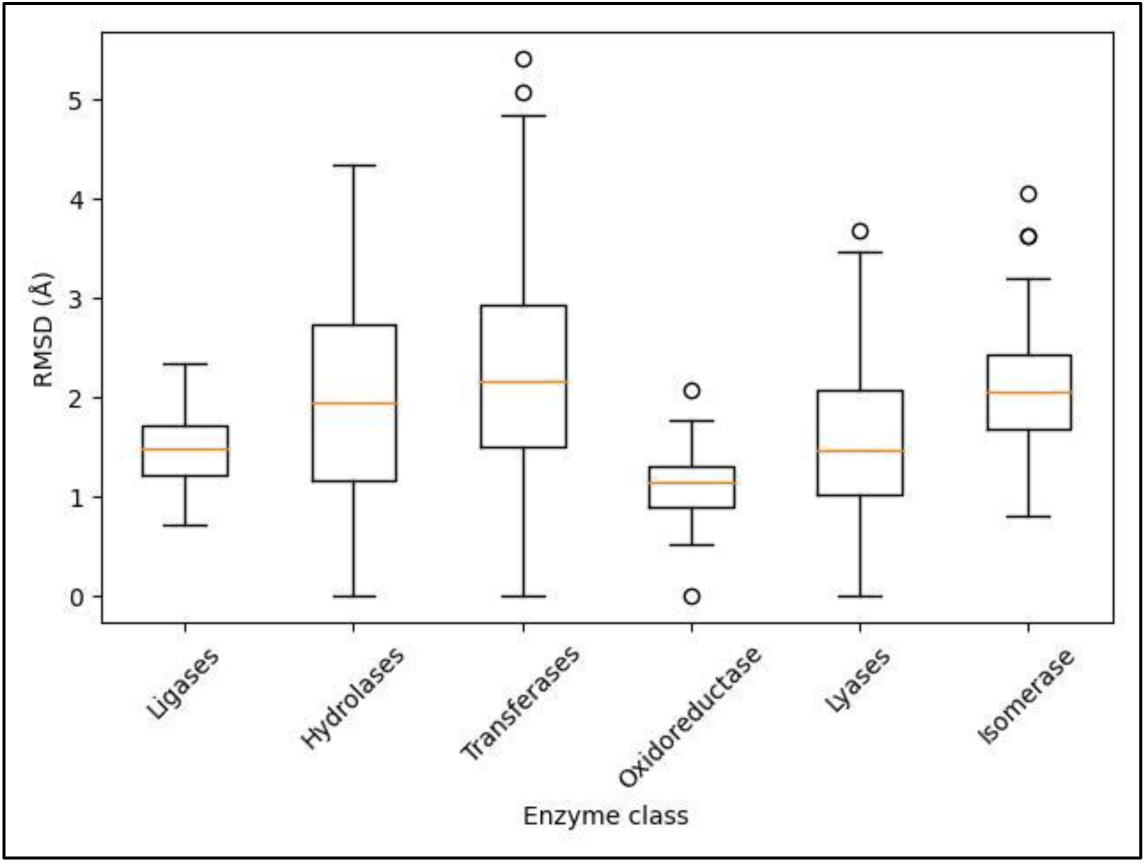
Performance of ligands for targets categorized as enzyme types in the validation set. While most enzyme types showed acceptable aggregate performance in terms of RMSD, transferases were observed to be a bit divergent. On the other hand, oxidoreductases showed exceptional performance.

**Figure 4:**
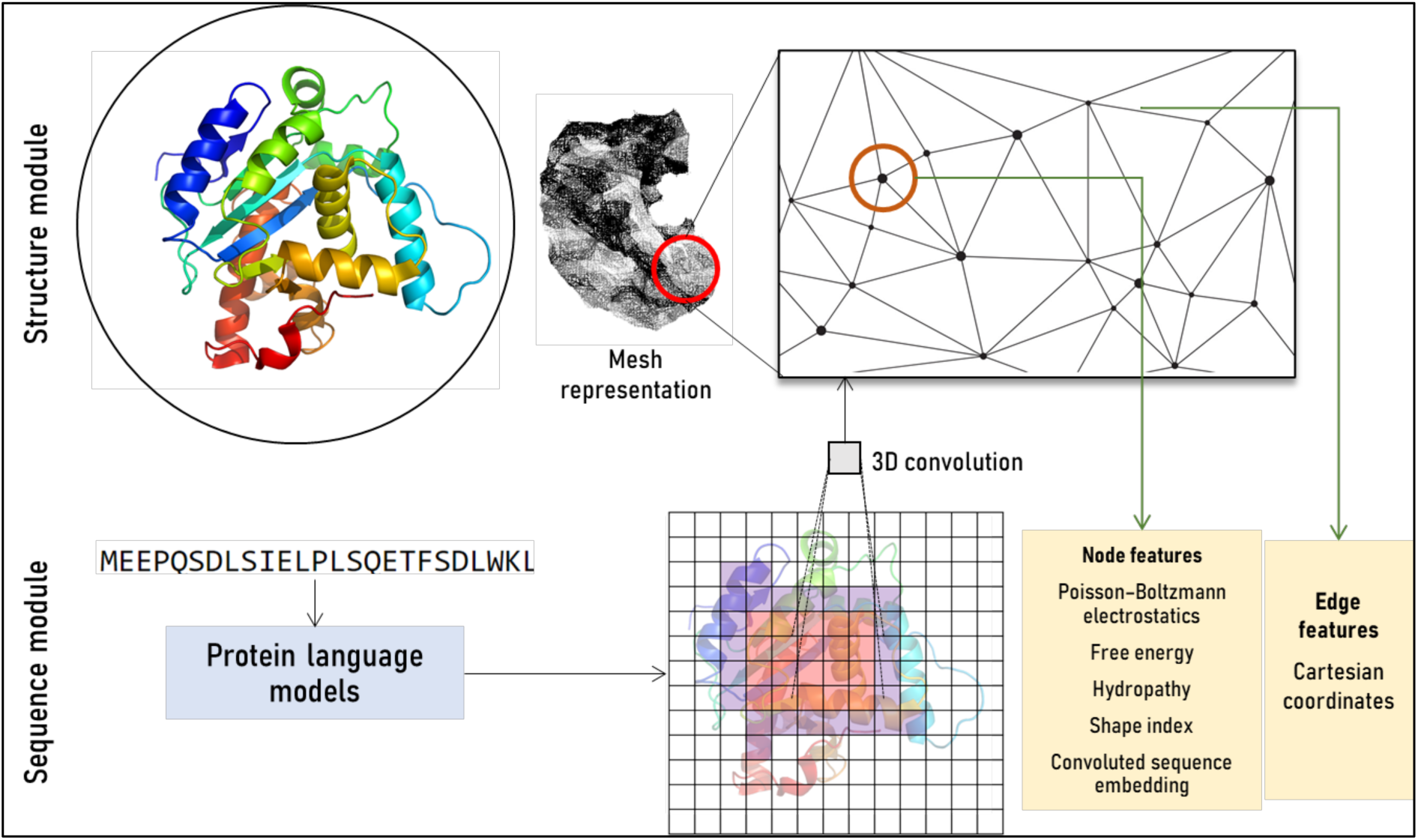
Featurization methodology for *Alphadock*. Separate structure and sequence modules were developed to process structural and sequence features individually. In order to integrate sequence embeddings to protein graphical structure, a convolution approach was employed. The extracted node and edge features are also compiled.

**Table 2:** Correlation between various factors for internal and external validation sets.

|  | <u>Variable 1</u> | <u>Variable 2</u> | <u>Correlation</u> |
| --- | --- | --- | --- |
| <u>Validation set</u> | <u>Score real</u> | <u>Score opt.</u> | <u>0.76</u> |
|  | <u>RMSD</u> | <u>Rot. bond</u> | <u>0.59</u> |
|  | <u>RMSD</u> | <u>No. atom</u> | <u>0.65</u> |
|  | <u>Score opt.</u> | <u>pK<sub>x</sub></u> | <u>0.48</u> |
| <u>CASP16 set</u> | <u>Score real</u> | <u>Score opt.</u> | <u>0.75</u> |
|  | <u>RMSD</u> | <u>Rot. bond</u> | <u>0.55</u> |
|  | <u>RMSD</u> | <u>No. atom</u> | <u>0.69</u> |
|  | <u>Score opt.</u> | <u>pK<sub>x</sub></u> | <u>0.52</u> |

For instance, protein size, location (cytoplasmic/membrane), function, structure and level of conservation within variants could affect the effectiveness of the graph representation. Since graph representation is a vital aspect of the workflow, it could have limited effectiveness in some edge cases. Although model performance is pretty acceptable for most of the cases in the validations set, the said limitation is particularly evident in the case of the transferase, where a number of outliers can be observed.

Going forward, it would also be very interesting to compare the effectiveness of other variations of MDNs, such as variational autoencoders, Generative Adversarial networks, and Normalizing Flow Models for solving the problem at hand. The proposed methodology and scoring function do suffer from some general limitations. For instance, the scoring function may struggle to account for dynamic aspects of ligand-receptor interactions, such as protein and ligand flexibility. The conformational changes that occur upon ligand binding and protein flexibility can significantly influence the binding affinity owing to the static nature of the available training data, and these complex phenomena are modelled in a simplified manner. Further, regarding the optimization method, since the differential evolution algorithm is a probabilistic algorithm, it does not guarantee that it will find the global minimum or maximum of the given function. It is also sensitive to the choice of parameters, such as the population size and the mutation and crossover probabilities, which can affect the performance of the algorithm.

In summary, this work introduces a novel workflow for modelling protein-ligand binding at a residue level by employing geometric deep learning. The method developed in this study differs from traditional scoring functions by using a deep neural network for learning a statistical potential that is based on the three-dimensional distribution of atoms. This allows the model to capture specific details of each protein-ligand complex individually. The learned potential, clubbed with an optimization algorithm, is then used to identify the optimal binding conformation, which minimizes the potential energy of the system. The results demonstrate that this approach is a reliable measure of binding capacity between a ligand and a protein, complementing the docking and virtual screening process. Moreover, the developed scoring system can be minimized with optimization algorithms like differential evolution or simulated annealing to arrive at the conformation having the highest propensity to interact with the given protein region. Further methodological developments in geometric deep learning and overall graph processing algorithms could significantly improve AI-assisted structure-based screening workflows to incorporate macromolecules of varying scale and complexity. Overall, this study reinforces the idea of representing molecular structures as graphs paired with sequence embeddings and the utility of geometric deep learning for estimating residue-level protein-ligand interactions.

## 4 Methods

### 4.1 Datasets and pre-processing

PDBbind database (v.2020) was employed to train the developed model [27]. It contained 19443 protein-ligand complexes along with their activity (in terms of disassociation constant and half-maximum inhibitory concentration). All complexes underwent standard processing steps like refinement and minimization. This was crucial for removing any unfavourable steric clashes or overlapping atoms in the initial complex, which can cause unrealistic distortions. These distortions can lead to inaccurate results and compromise the reliability of the downstream analysis. Moreover, the spatial coordinates of protein and ligand were ideally required in their minimum energy state so as to generate reliable ground truths (pairwise distances between each ligand-target node). Glide package from the Schrodinger Suites program was employed to preprocess, refine and minimize the complexes [28]. All codes that were developed for the overall pipeline are available on GitHub (https://github.com/TeamSundar/alphadock)

### 4.2 Representing protein and ligand structures as graphs

A data processing pipeline, based on existing methods was developed to represent the 3D structure of the protein in a trainable form. The underlying idea was to represent topological characteristics of the binding region in a graphical structure. An overview of the various modules employed are summarized in **Error! Reference source not found.**

The given regions on the protein surface (designated by taking the bound ligand molecule as a reference) were triangulated using MSMS that resulted in a graph structure, generally denoted as a mesh [29]. A probe radius of 1.5 Å and a node density of 3.0 nodes per Å^2^ was for generating the mesh, which was followed by further processing on PyMesh. Adaptive Poisson-Boltzmann Solver (APBS) was employed to calculate the surface characteristics as features for the graph embeddings. Poisson-Boltzmann continuum electrostatics, free electron/proton donors, hydropathy and shape index were calculated for the given binding region [30]. A complete list of features is compiled in **Table 3**.

**Table 3:** List of features that were embedded as node features in the graph representations of protein and drugs.

|  |  |
| --- | --- |
| Ligand features | Single, Double, Triple, Aromatic<br>As, Be, Br, C, Ce, Co, Cu, F, Fe, Hg, I, Ir, Mg, N, O, Os, P, Pt, Re,<br>Ru, Rh, S, Sb, Se, Si, V, Zn |
| Protein Features | Poisson-Boltzmann continuum electrostatics<br>Free electron/proton donors<br>Shape index<br>Hydropathy<br>Convolved sequence embeddings |

This pipeline resulted in a mesh structure *G*^t^ = (*N*^t^, *ℰ*^t^), representing the protein surface. Each node 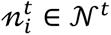 was populated with a four-dimensional vector containing the features calculated by APBS. Similarly, each edge 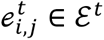 encoded the spatial coordinates of nodes. It should be emphasized that only the regions that were in vicinity of the bound ligand were used to construct these graph structures in the interest of computational complexity and hardware limitations. Encoding ligand molecules was relatively straightforward, wherein they were represented as undirected graphs *G*^1^ = (*N*^1^, *ℰ*^1^). Atoms and bonds were denoted by 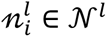 (28D one hot vectors for element as node features) and 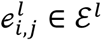 (4D one-hot vectors for bond-type as edge features) respectively. While no information regarding the molecular conformation was included in the encodings, it was expected from the model to perceive the correct conformation from the distance information embedded in the generated graphs. A general processing and architectural setup have been summarized in Error! Reference source not found.

### 4.3 Graph neural networks and model architecture

Graph neural networks (GNNs) are based on the idea of propagating information through a graph by updating the representations of the nodes based on their neighboring nodes. GNNs typically consist of multiple layers, with each layer processing the representations of the nodes and edges in the graph and producing updated representations for the next layer. The process of updating the representations at each layer is called “message passing,” and it is typically done using a variant of the following equation:

Messaging:

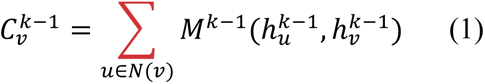

Readout:

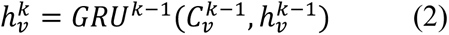

Where eq (1) articulates the information exchange between nodes in a graph. At each iteration or layer *k*, each node *v* aggregates information from its neighbors (*N*(*v*)) by applying a message function *M*^(*k*-1)^ to the hidden states of both the node itself (*h_v_*^(*k*-1)^) and its neighbors (*h_u_*^(*k*-1)^)The result, denoted as (*C_v_*^(*k*-1)^), represents the accumulated information that node *v* receives from its local neighborhood. This step allows the network to consider the context and relationships of a node within the graph. Furthermore, the readout equation (2), outlines how the hidden state of a node *v* is updated based on the aggregated messages obtained in the Message Passing step. A Gated Recurrent Unit (GRU) operation, denoted as *GRU*^(*k*-1)^ takes as input both the aggregated messages (*C_v_*^(*k*-1)^) and the previous hidden state *h*^(*k*-1)^) The GRU is a type of recurrent neural network (RNN) that selectively incorporates information from both sources, allowing the model to capture temporal dependencies and update the hidden state for the current layer 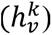. This iterative process is typically repeated across multiple layers in the GNN, enabling the network to capture intricate patterns and relationships within the graph-structured data. In essence, the Message Passing and Readout operations enable the GNN to iteratively refine its understanding of the graph, incorporating information from neighboring nodes and updating node representations over multiple layers.

Ligand and target graphs were passed through two independent residual GNNs to extract features. Nodes and edges were updated based on their neighboring nodes and edge types. Ultimately, the updated node features for ligands and targets were concatenated pairwise. Processed graph data from GNNs were passed through MDNs, which are a type of neural network architecture that is commonly used in probabilistic modeling tasks, such as modeling complex data distributions or generating novel samples from a learned distribution. MDNs model the conditional probability distribution of a target variable (e.g., distance between any drug-target node pair) as a mixture of several Gaussian distributions, where each Gaussian represents a different mode of distribution.

The implementation had two parts: a standard neural network that maps an input to a set of parameters for the Gaussian mixture distribution (modelled by MLP to create a latent representation *h_i_*_,*j*_ of concatenated target and ligand features, where *i* and *j* are the indices of each target and ligand node respectively), and a set of mixture density components, each of which is defined by a mean vector, covariance matrix, and weight (or mixture coefficient). During training, the parameters of the MDN were optimized to maximize the likelihood of the training data given the mixture model. In other words, the network learned to generate predictions that are consistent with the observed pairwise atomic distances, while also being able to capture the complex structure of the underlying data distribution.

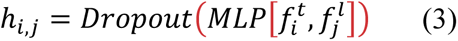

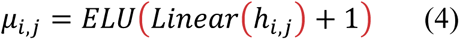

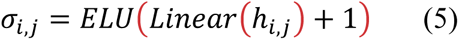

### 4.4 Training

The ***Alphadock*** model was trained to minimize loss function depicted in eq. 4-6 where *l_MDN_*, corresponds to the mixture density network loss, while *l*a*_tom_* and *l_bond_* corresponds to cost associated with atom and bond types. Adam optimizer was used along an adaptive learning rate of 0.0001 for updating model weights.

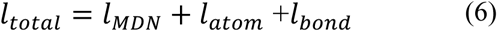

It should be noted that *l_MDN_*, corresponds to the negative log likelihood of the distance separating a particular ligand-target nodes pair.

Also, the given loss function can be used to derive a scoring potential *U*_7_ that would be specific to a particular ligand-protein complex.

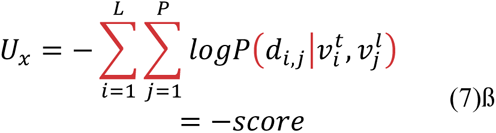

The model was trained for 200 epochs with a batch size of 32 protein-ligand pairs. As described earlier, the network was trained to estimate the mixture model parametrized by mean, variances and mixture coefficients. The network was trained for a varied number of gaussians to identify the optimal structure of the mixture model.

### 4.5 Benchmarking

The effectiveness of the method developed in this study was evaluated on the CASF benchmark (2016). It hosts a total of 285 protein-ligand complexes with a host of decoys. All the complexes were processed as described earlier. The developed method was evaluated under three metrics, namely, docking power, screening power and reverse screening power. Docking power is the potential of a scoring function to correctly rank true ligand among a host of decoys for a given protein target. In case of screening power, it gauges the capability of the scoring function to identify the correct ligand among random molecules. Similarly, reverse screening power determines the ability of the scoring function to correctly identify the true binding target for a given ligand among multiple random proteins.

### 4.6 Predicting binding conformers

Differential evolution was employed to optimize the conformation of the ligand molecule. For a given ligand molecule, Vector of Euler angles, dihedral angles of all rotatable bonds and relative positioning of the ligand in the Euclidean space were iteratively optimized so as to minimize the potential *U*_7_ defined in eq. 4-7. Optimization was run for 500 iterations with a variable mutation constant at each generation. Further, Euler angles for the rotatable bonds were restricted from -π to π.

## Notes

https://github.com/TeamSundar/alphadock

### Competing Interest Statement

The authors have declared no competing interest.

### Summary of Updates

Table 2 has been added to the revised manuscript to emphasize the correlation abound various parameters. Equations 1 and 2 in Section 4.3 (Graph neural networks and model architecture) have now been explained in detail for clarity.

